# A novel compound that disrupts mitotic spindle poles in human cells

**DOI:** 10.1101/2020.08.14.251058

**Authors:** Dilan Jaunky, Mathieu Husser, Kevin Larocque, Peter Liu, Sajinth Thampipillai, Pat Forgione, Alisa Piekny

**Author notes:** Equal contribution (Mathieu Husser, Kevin Larocque). Correspondence: Alisa Piekny, PhD, Associate Professor.

## Abstract

We characterize the mechanism of action of a new microtubule-targeting compound in cells. Microtubule-targeting drugs are used as successful anti-cancer therapies. We synthesized a family of compounds that share a common scaffold and have several functional groups amenable to modifications. We found that one of the active derivatives, C75, reduces cell viability and prevents microtubule polymerization *in vitro*. In this study, we explore the phenotypes caused by C75 in cells. It causes mitotic arrest and spindle phenotypes in several cancer cell lines in the nanomolar range. C75 can bind to the Colchicine-pocket on tubulin *in vitro*, but causes different effects on microtubules in cells. While Colchicine causes a decrease in microtubules and spindle pole collapse without re-growth, similar concentrations of C75 cause a rapid loss of microtubules and spindle pole fragmentation followed by microtubule re-growth to form multipolar spindles. In addition, C75 and Colchicine synergize for reduced viability and spindle phenotypes. Importantly, the phenotypes caused by C75 are similar to those caused by the depletion of ch-TOG, a microtubule polymerase, and tubulin and ch-TOG are displaced and oscillate in C75-treated cells. This suggests that C75 causes microtubule depolymerization in cells either directly or indirectly via inhibiting ch-TOG. This unique effect of C75 on microtubules warrants further exploration of its anti-cancer potential.

## Introduction

Dynamic microtubules are required for mitotic spindle assembly and function, and drugs that suppress these dynamics are used to treat cancer (Cirillo et al. 2017; Kozielski 2015; Musacchio 2015). At the G2/M transition, there is an increase in microtubule growth from the maturing centrosomes. Dimers of α-tubulin and β-tubulin form 13 protofilaments that roll into a tubule, which are templated by the γ-tubulin ring complex (γ-TURC; Oakley et al. 2015; Martin and Akhmanova 2018). GTP-binding and hydrolysis influences microtubule structure and can promote growth or catastrophe (Wordeman 2019; Martin and Akhmanova 2018; Brouhard and Rice 2018). Straightening of the protofilament plus-ends are crucial for microtubule stability and growth. During catastrophe or under drug-induced conditions that destabilize the plus-end, the protofilaments will adopt a curved state and bend away from the lumen (Gigant et al. 2005; Akhmanova and Steinmetz 2015; Goodson and Jonasson 2018). The growth and shrinkage of microtubules is required to form stable kinetochore attachments with sister chromatids, and many microtubule-associated proteins (MAPs) influence microtubule dynamics by stabilizing the minus and/or plus ends (Goodson and Jonasson 2018). MAPs include microtubule motors that bundle and organize microtubules to form bipolar spindles, as well as enzymes such as ch-TOG and MCAK that control microtubule polymerization and depolymerization, respectively (Widlund et al. 2011; Zhang et al. Walczak 2008; Gergely et al. 2003; Holmfeldt et al. 2004; Barr and Gergely 2008; Brouhard et al. 2008; Barr and Bakal 2015; Brouhard and Rice 2018). When stable kinetochore attachments are achieved by both sister chromatids, this generates tension that is sensed by the spindle assembly checkpoint for mitotic exit and the segregation of sister chromatids (Musacchio 2015). Microtubule-targeting drugs that suppress microtubule dynamics and prevent the formation of stable kinetochore attachments can lead to cell cycle arrest or catastrophe, and cell death.

Several different microtubule-targeting drugs are currently being used to treat cancers. One of the hallmarks of cancer is uncontrolled cell proliferation, and this property can make cancer cells more responsive to compounds that arrest different stages of the cell cycle compared to healthy cells (Hanahan and Weinberg 2011; Dominguez-Brauer et al. 2015). Taxol™ is a microtubule-targeting compound that causes mitotic arrest or catastrophe and is used to treat a plethora of cancers (Weaver 2014). Taxol binds to microtubules and stabilizes the lattice to prevent depolymerization (Kumar 1981; Parness and Horwitz 1981). This prevents the formation of stable kinetochore attachments, and failed segregation of sister chromatids (Zasadil et al. 2014). Depending on the other genetic changes in the cell, this can trigger the spindle assembly checkpoint, or cause mitotic catastrophe (Zasadil et al. 2014). Although Taxol offers some success in the clinic, it can cause severe side-effects and some patients develop resistance (Kavallaris 2010). Thus, there is a need to develop alternative drugs that can reduce the concentrations of Taxol needed for treatment to reduce side-effects and resistance.

Cancer cells often accumulate mutations that alter the expression and/or function of MAPs, motor proteins and microtubule-regulating enzymes, which causes them to have increased sensitivity to microtubule-targeting drugs (Cirillo et al. 2017; Kozielski 2015; Musacchio 2015; Brito and Rieder 2010). Coupling the depletion of MCAK and treatment with low concentrations of Taxol in cancer cells causes a dramatic increase in the proportion of cells in mitosis with multipolar spindles, and increased apoptosis (Hedrick et al., 2008). In colorectal cancer cells, Aurora A kinase, which regulates centrosome maturation and plus-end dynamics for kinetochore attachments, and ch-TOG (CKAP5), a microtubule polymerase, are over-expressed (Holmfeldt et al. 2004; Fielding et al. 2011; Barr and Bakal 2015; Al-Bassam and Chang 2011; Byrnes and Slep 2017; Yu et al. 2016; Hood et al. 2013). Their overexpression correlates with increased rates of microtubule assembly and chromosomal instability (CIN) due to excess lagging chromosomes (Ertych et al., 2014, Byrnes and Slep 2017). These properties could make mitotic spindles sensitive to further disruption by microtubule-targeting drugs compared to healthy cells. However, if CIN cells have lost tumor suppressors or checkpoint regulators, they could lose their responsiveness to these drugs and retain tumor growth properties (Ertych et al., 2014).

Another class of microtubule targeting compounds prevents polymerization or causes depolymerization of microtubules depending on their concentration and accessibility. These altered dynamics cause mitotic arrest or catastrophe similar to Taxol and have been used or considered for use as anti-cancer therapies. Colchicine was discovered by Gary Borisy (1967) and led to the biochemical purification of tubulin (Borisy and Taylor 1967). Colchicine binds to a deep pocket on β-tubulin and prevents growth or causes catastrophe at the plus-end (Fitzgerald 1976; Kumar et al. 2016; Massarotti et al. 2012; Wang et al. 2016). Colchicine is used in the clinic to treat gout, but failed as an anti-cancer therapy (Field et al. 2015; Jordan and Kamath 2007). Vinblastine also prevents microtubule polymerization or causes microtubule depolymerization, and is being used as an anti-cancer drug (Martino et al. 2018; Gigant et al. 2005). It binds to the interface of the heterodimer and introduces a molecular wedge causing the protofilaments to adopt a curled conformation that effectively destabilizes the microtubule (Martino et al. 2018; Gigant et al. 2005; Jordan and Kamath 2007). Vinblastine causes lower side-effects compared to Colchicine, and this higher tolerance could explain its success as an anti-cancer drug compared to Colchicine. In addition to having distinct binding sites, they could have different properties *in vitro* and *in vivo*, including accessibility to the minus vs. plus ends that alters their affinity and how they affect the microtubule polymer. *In vitro*, Vinblastine favours depolymerization at the minus ends of polymerized microtubules and suppression of dynamics at the plus ends (Panda et al., 1996). In HeLa cells, low concentrations (*e*.*g*. 2 nM) of Vinblastine did not cause obvious changes in the microtubule polymer mass, but blocked mitosis by decreasing kinetochore attachments, as well as the dissociation of mother and daughter centrioles and loss of centriole ultrastructure (Jordan et al., 1992; Wendell et al., 1993).

We recently synthesized a novel family of compounds that target microtubules (*manuscript submitted*). Our approach was to strategically design and synthesize a family of compounds with properties ideal for *in vivo* use and screen derivatives for their ability to arrest cancer cell proliferation. The compounds share a common core thienoisoquinoline scaffold amenable to modifications via structure-activity-relationship (SAR) studies (Chen et al. 2014). We identified several derivatives that cause mitotic arrest with an IC_50_ for viability in the nanomolar range. Importantly, some derivatives have no or little effect on cells (*e*.*g*. C87, IC_50_ >10 uM) while others have high efficacy (*e*.*g*. C75, IC_50_ between 0.1 – 0.4 uM depending on the cell line). We also found that C75 prevents microtubule polymerization *in vitro*, suggesting that it can bind to tubulin.

Here, we characterize the spindle phenotypes caused by C75 in cells to elucidate its mechanism of action in cells. We found that despite being able to bind to the same pocket as Colchicine *in vitro*, C75 has different effects on microtubule polymers and mitotic spindle organization. C75 causes cancer cells to arrest in mitosis with aberrant spindles, misaligned chromosomes and spindle pole fragmentation. Adding C75 and Colchicine together showed synergistic effects at several concentrations in HeLa and HCT 116 cells, suggesting differences in their binding properties or accessibility. In live cells, microtubules are lost with C75-treatment, but unlike Colchicine, C75 also causes spindle pole fragmentation and microtubule re-growth to form multipolar spindles. The spindle phenotypes that arise due to C75 are similar to those caused by ch-TOG RNAi, suggesting that C75 promotes microtubule depolymerization in cells. Interestingly, spindle pole components are displaced and oscillate upon C75-treatment, suggesting that it has accessibility to the minus ends. Therefore, C75 is a novel microtubule-targeting drug with effects on microtubule polymers that appear to be distinct from those caused by Colchicine. While it promotes depolymerization, the re-growth and formation of multipolar spindles make it an interesting compound to explore further for *in vivo* use to disrupt tumor growth.

## Material and Methods

### Cell culture and drug treatments

HeLa and HFF-1 cells were grown in DMEM (Wisent), while A549 cells were grown in F12K media (Wisent) and HCT 116 (p53-/-) cells were grown in McCoy’s media (Wisent) in humidified incubators at 37°C with 5% CO_2_. All media were supplemented with 10% fetal bovine serum (FBS; Thermo Scientific), 2 mM L-glutamine (Wisent), 100 U penicillin (Wisent), and 0.1 mg/mL streptomycin (Wisent).

C87 and C75 were stored at -20°C as 1 mM stocks in dimethyl sulfoxide (DMSO). Working stocks of 100 *μ*M C75 or C87 were made in DMSO:H_2_O (9:1). Colchicine (Sigma) was dissolved in ethanol as a 1 M stock and diluted to 10 *μ*M before use. Final concentrations of DMSO or ethanol were kept below 0.5%.

### Microtubule assays

Microtubule polymerization assays were performed using 1.5 mg/mL purified tubulin (Cytoskeleton, Inc) that was taken from a 10 mg/mL flash frozen stock, thawed on ice and diluted in G-PEM buffer (80 mM PIPES pH 6.9, 2 mM MgCl_2_, 0.5 mM EGTA and 1 mM GTP with 20% glycerol) with 10% DMSO (control) or 250 nM C75. Assays were performed in a preheated 50 uL sub micro Z15 black Q/Spectrosil cuvette in a Varian Cary 1 spectrophotometer. Reagents were added to the cuvette and blanked immediately before recording absorbance at 340 nm in 0.1 second intervals for 45 minutes.

We determined the effect of C75 on polymerized microtubules by sedimentation. Lyophilized microtubules (Cytoskeleton, Inc) were reconstituted to 5 mg/mL in PM buffer (15 mM PIPES pH 7.0, 1 mM MgCl_2_, and 2 mM Taxol) and aliquots were snap frozen in liquid nitrogen and stored at -80°C. Aliquots were thawed on ice in a circulating water bath, then diluted to 2 mg/mL in PM buffer at room temperature. Microtubules (1 μM) were incubated with 5 μM of C75, Taxol, or Colchicine in a general tubulin buffer (80 mM PIPES pH 6.9, 2 mM MgCl_2_, 0.5 mM EGTA) for 15 minutes. Then the samples were centrifuged at 16,000 xg for 1 hour. The supernatant and pellets were resuspended in sample buffer without SDS or DTT and run on by native PAGE on an 8% polyacrylamide gel.

Colchicine-competition assays were performed with 2 μM purified tubulin (Cytoskeleton, Inc) taken from a 10 mg/mL stock prepared in tubulin buffer as above, and the binding assays were performed with 1 μM Colchicine or 1 μM Colchicine with 500 nM C75 in 25 mM PIPES buffer, pH 6.8 in a 600 μL Q fluorometer cell, Z=20 (Varian) using a fluorescence spectrophotometer Cary Eclipse (Varian) at 25 °C for 3 hours. Reaction mixtures were excited at 350 nm and the emission was measured from 380 to 500 nm (Bhattacharyya and Wolff, 1974). Emission values were collected at 30-minute intervals and normalized to the spectra recorded at time zero. All values were exported as excel files and graphs were generated in Prism V8.1.0 Graphpad.

### Viability assays

Assays were performed to determine the viability of cells after treatment with C75 and/or Colchicine. The concentration of Colchicine or C75 used in the combination experiments was the highest concentration that caused little to no toxicity based on dose-response curves. HFF-1, HeLa, A549 and HCT 116 cells were plated with 4,000 cells/well in 96-well dishes and left overnight to adhere. Drug dilutions were prepared and added to the cells using an acoustic liquid handler (LabCyte ECHO 550). After 3 population doubling times, cells were assessed for viability using the WST-8 cell proliferation assay kit (Cayman Chemicals). Absorbance readings at 450 nm were collected using the TECAN 200 PRO plate reader. Values for each replicate were adjusted to the controls and plotted using GraphPad Prism 7 to generate graphs and IC_50_ values. All assays were performed in triplicate for each treatment. For the combination assays, C75 or Colchicine were repeated alongside the combination treatments to ensure accurate comparison.

### Flow cytometry

Flow cytometry was used to measure changes in the proportion of cells in different stages of the cell cycle after C75-treatment. HeLa, HCT 116 and A549 cells were grown to 80% confluency and treated with a range of C75 concentrations for 8 hours. Cells were harvested in falcon tubes, then fixed with 70% cold ethanol and washed with cold phosphate buffered saline (PBS; Wisent). Cells were permeabilized and stained for 15 minutes at 37 °C with a solution containing 500 µg/mL PI (Sigma) in PBS with 0.1% (v/v) Triton X-100 and DNAse-free RNAse A (Sigma). Cells were protected from light and measured for PI intensity using the BD LSRFortessa flow cytometer with excitation at 561 nm and detection at 600 nm (LP) using the D-BP filter. Each treatment was done in triplicate with 20,000 cells analyzed per sample. Data was exported and plotted using GraphPad Prism 7 to make bar graphs.

### Immunofluorescence staining

Immunofluorescence was performed to monitor mitotic spindle phenotypes. Cultured cells were plated on coverslips at a confluency of 40-50% and left overnight to adhere. Cells were fixed using freshly prepared ice-cold 10% w/v cold trichloroacetic acid (TCA) for 14 minutes at 4°C. Cells were washed with PBST (0.3% Triton X-100) and kept at 4°C prior to staining. After blocking, fixed cells were immunostained for microtubules using 1:400 mouse anti-α-tubulin antibodies (DM1A; Sigma) or centrosomes using 1:400 mouse anti-γ-tubulin antibodies (Santa Cruz Biotechnology) or mouse anti-Centrin 2 (clone 20H5; Sigma), and centromeres using 1:500 human anti-centromere antibodies (ACA; Sigma) for 2 hours at room temperature. After washing, anti-mouse Alexa 488 and anti-human Alexa 647 (Invitrogen) secondary antibodies were used at a 1:500 dilutions for 2 hours at room temperature. After washing, 4′,6-Diamidino-2′-phenylindole dihydrochloride (DAPI; Sigma) was added for 5 minutes. Cells were then washed with PBST, followed by a wash with 0.1 M Tris pH 9, then a drop of mounting media (0.5% propyl gallate in 50% glycerol) was added to the coverslip, which was mounted onto a slide and sealed.

### Microscopy

Fixed slides were imaged using the Nikon-TIE inverted epifluorescence microscope with Lambda XL LED light sources, using the 60x/1.4 oil objective, a Piezo Z stage (ASI), a Photometrics Evolve 512 EMCCD camera and Elements acquisition software (Nikon). Exposures were determined by control cells, and the same settings were used in the treatment conditions. Images were acquired as 1 µm Z-stacks and exported as TIFFs, which were opened as maximum intensity Z-stack projections in Image J (NIH). Merged colour images were converted into 8-bit images and imported into Illustrator (Adobe) to make figures.

For live imaging, HeLa cells were plated to 50-60% confluency on 25 mm round coverslips (No. 1.5; Neuvitro). Cells were treated with 75 nM Hoescht 33342 and 200 nM SiR-tubulin (Cytoskeleton Inc.) for 90 minutes prior to imaging. Depending on the experiment, HeLa cells were transfected with a plasmid that expresses RNAi-resistant ch-TOG:GFP plus a hairpin RNA (sh ch-TOG) to knockdown endogenous protein and minimize overexpression (Addgene ID# 69113). HeLa cells stably expressing GFP:tubulin were previously generated (van Oostende Triplet et al., 2014). Coverslips were transferred to a 35 mm chamlide magnetic chamber (Quorum) and kept at 37 °C with 5% CO_2_ using an INU-TiZ-F1 chamber (MadCityLabs). Images were acquired using the 60x/1.4 oil objective on the Nikon Livescan sweptfield confocal microscope with an Andor iXon X3 EMCCD camera and Elements acquisition software (Nikon). The 405 nm, 480 nm and 640 nm lasers were used to image Hoescht, GFP and sir-Tubulin, respectively, with a quad filter and exposures set to controls. Z-stacks of 1 µm were collected using a NI-DAQ piezo Z stage (National Instruments). Images were used to make figures as described above.

To analyze the recovery of microtubules after cold-treatment in cells, HeLa cells were plated onto coverslips and transferred to a 35 mm chamlide magnetic chamber (Quorum), which was placed in an ice-cold water bath to cause spindle collapse. After 30 minutes, the cells were transferred to 37 °C with 5% CO_2_ using an INU-TiZ-F1 chamber (MadCityLabs) and C75 or Colchicine was added after acquisition of the first timepoint. Images were acquired using the 60x/1.4 oil objective on the Nikon Livescan sweptfield confocal microscope with an Andor iXon X3 EMCCD camera and Elements acquisition software (Nikon). The 405 nm and 480 nm lasers were used to image Hoescht 33342, GFP:tubulin respectively, with a quad filter and exposures set to controls. Z-stacks of 1 µm were collected using a NI-DAQ piezo Z stage (National Instruments). Images were used to make figures as described above.

### Analysis

For the analysis of phenotypes in the Figures 2, 3, S3, 4 and 6, either whole fields of view (Figure 2A) or z-stack projections of individual cells were used (Figures 2B, C, D, 3B, C, S3B, 4B, 6A, B). To calculate the proportion of mitotic cells based on cell rounding (Figure 2A) we counted the following number of HeLa cells on average: 0 nM n=568, 100 nM n=847, 200 nM n=266, 300 nM n=105, 400 nM n=96, 500 nM n=75), A549 cells: 0 nM n=461, 100 nM n=519, 200 nM n=181, 300 nM n=154, 400 nM n=150, 500 nM n=253), and HCT 116 cells: 0 nM n=813, 100 nM n=742, 200 nM n=297, 300 nM n=296, 400 nM n=314, 500 nM n=301 (N=4 experimental replicates). To calculate the proportion of cells with different mitotic spindle phenotypes (Figure 2B and C), the following number of HFF-1 cells were counted on average after treatment with DMSO n=28, 300 nM C87 n=24 and 300 nM C75 n=13, HeLa cells after treatment with DMSO n=45, 300 nM C87 n=36 and 300 nM C75 n=109, A549 cells with DMSO n=29, 300 nM C87 n=32 and 300 nM C75 n=63, and HCT 116 cells with DMSO n=41, 300 nM C87 n=32 and 300 nM C75 n=66 (N=3 experimental replicates). Fewer HFF-1 cells were analyzed compared to the other cell lines due the low proportion of mitotic cells in the population. To determine the number of centrin-2-positive foci in HeLa cells (Figure 2D), an average of 25 cells were counted for the control (DMSO-treated) and 30 cells after treatment with 300 nM C75 (N=3 experimental replicates). To calculate changes in the proportion of mitotic spindle phenotypes in HeLa cells after treatment with C75 and/or Colchicine (Figures 3B and C), we counted the following average numbers of cells: DMSO n=51, 20 nM Colchicine n=62, 50 nM Colchicine n=72, 100 nM Colchicine n=78, 100 nM C75 n=42, 200 nM C75 n=46, 300 nM C75 n=59, 400 nM C75 n=69, 500 nM C75 n=65, 100 nM C75 + 20 nM Colchicine n=70, 200 nM C75 + 20 nM Colchicine n=89, 300 nM C75 + 20 nM Colchicine n=94, 400 nM C75 + 20 nM Colchicine n=98 and 500 nM C75 + 20 nM Colchicine n=83 (N=3 experimental replicates). To calculate changes in the proportion of mitotic spindle phenotypes in HCT 116 cells after treatment with C75 and/or Colchicine as above (Figure S3B), we counted the following average number of cells: DMSO n=44, 20 nM Colchicine n=39, 50 nM Colchicine n=54, 100 nM Colchicine n=51, 100 nM C75 n=42, 200 nM C75 n=40, 300 nM C75 n=82, 400 nM C75 n=56, 500 nM C75 n=65, 100 nM C75 + 20 nM Colchicine n=43, 200 nM C75 + 20 nM Colchicine n=52, 300 nM C75 + 20 nM Colchicine n=69, 400 nM C75 + 20 nM Colchicine n=76 and 500 nM C75 + 20 nM Colchicine n=81 (N=3 experimental replicates). To count the proportion of cells with multipolar or bipolar spindles in HeLa cells after release from C75 or Colchicine (Figure 4B), we counted the following number of average cells: DMSO n=61, 500 nM C75 n=50 and 500 nM Colchicine n=64 (N=3 experimental replicates). To determine the proportion of cells with mitotic spindle phenotypes and compare these to ch-TOG RNAi treatment (Figure 6A and B), we counted the following number of average cells: DMSO n=81, 200 nM C75 n=242, 300 nM C75 n=143, ch-TOG RNAi n=534 (N=3 experimental replicates).

**Figure 1.**
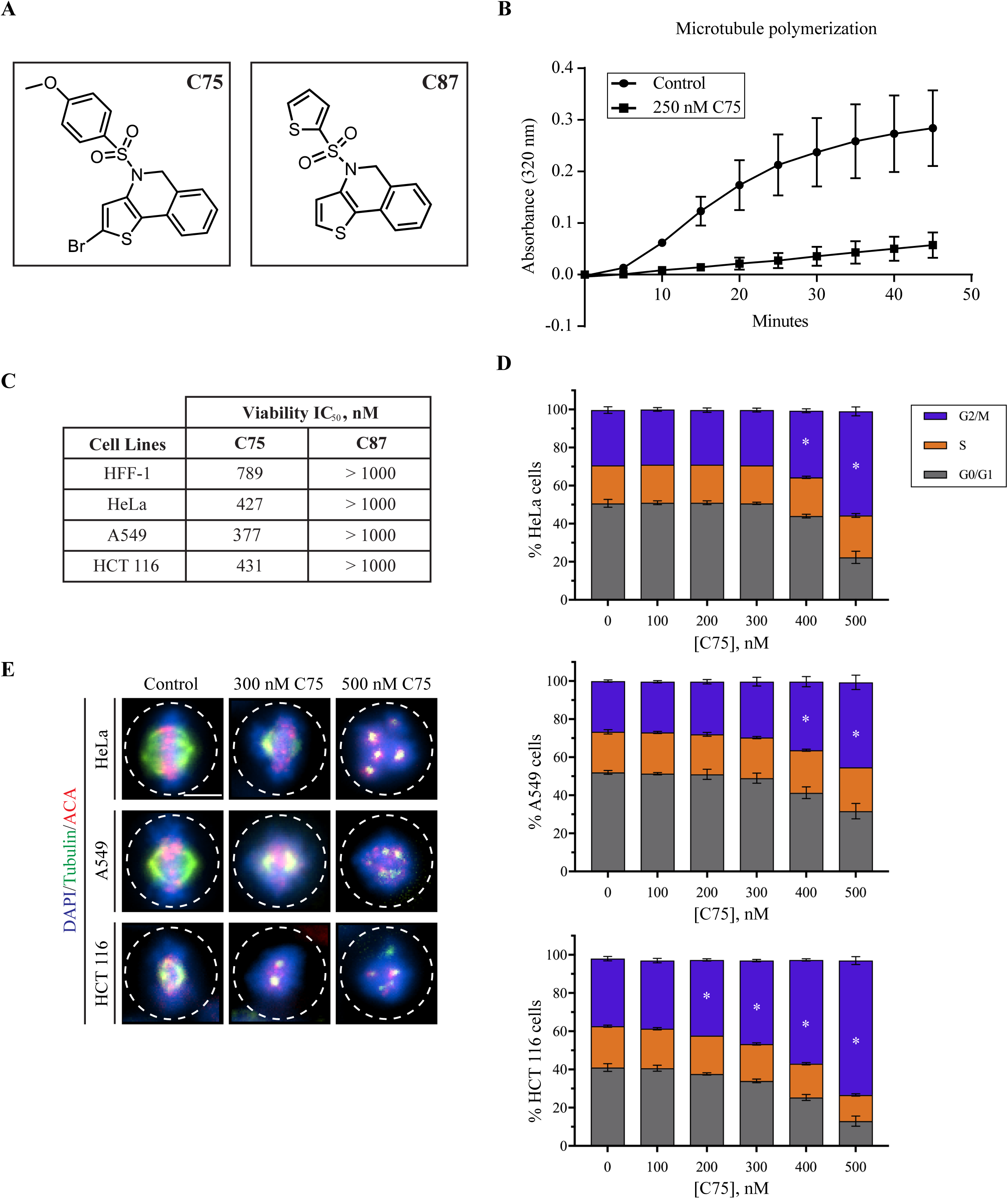
C75 is a thienoisoquinoline compound that causes G2/M arrest in cultured human cells. **A)** The structures of C75 and C87, a derivative with minimal activity, are shown. **B)** A graph shows the polymerization of microtubules *in vitro*, measured by changes in absorbance at 320 nm (X-axis) over time (minutes; Y-axis). Tubulin was polymerized in the presence of 250 nM C75 or DMSO (control). **C)** A table shows the IC_50_ for the viability of HFF-1, HeLa, A549 and HCT 116 over three population doubling times after treatment with C75 or C87 (N=3). **D)** Bar graphs show the distribution of cells in G0/G1, S and G2/M phases of the cell cycle measured by flow cytometry, for HeLa, A549 and HCT116 cells treated with increasing concentrations of C75 for 8 hours (n = 20,000 cells per treatment; N=3 experimental replicates). Asterisks indicate statistical significance using two-way ANOVA test and post-hoc Tukey’s multiple comparison test with a 95% CI, multiplicity adjusted *p* =< 0.0004. **E)** Images show fixed HeLa, A549 and HCT116 cells immunostained for DNA (DAPI; blue), tubulin (green), and centromeres (ACA; red) after treatment with 300 or 500 nM of C75 for 8 hours. The scale bar is 10 μm.

**Figure 2.**
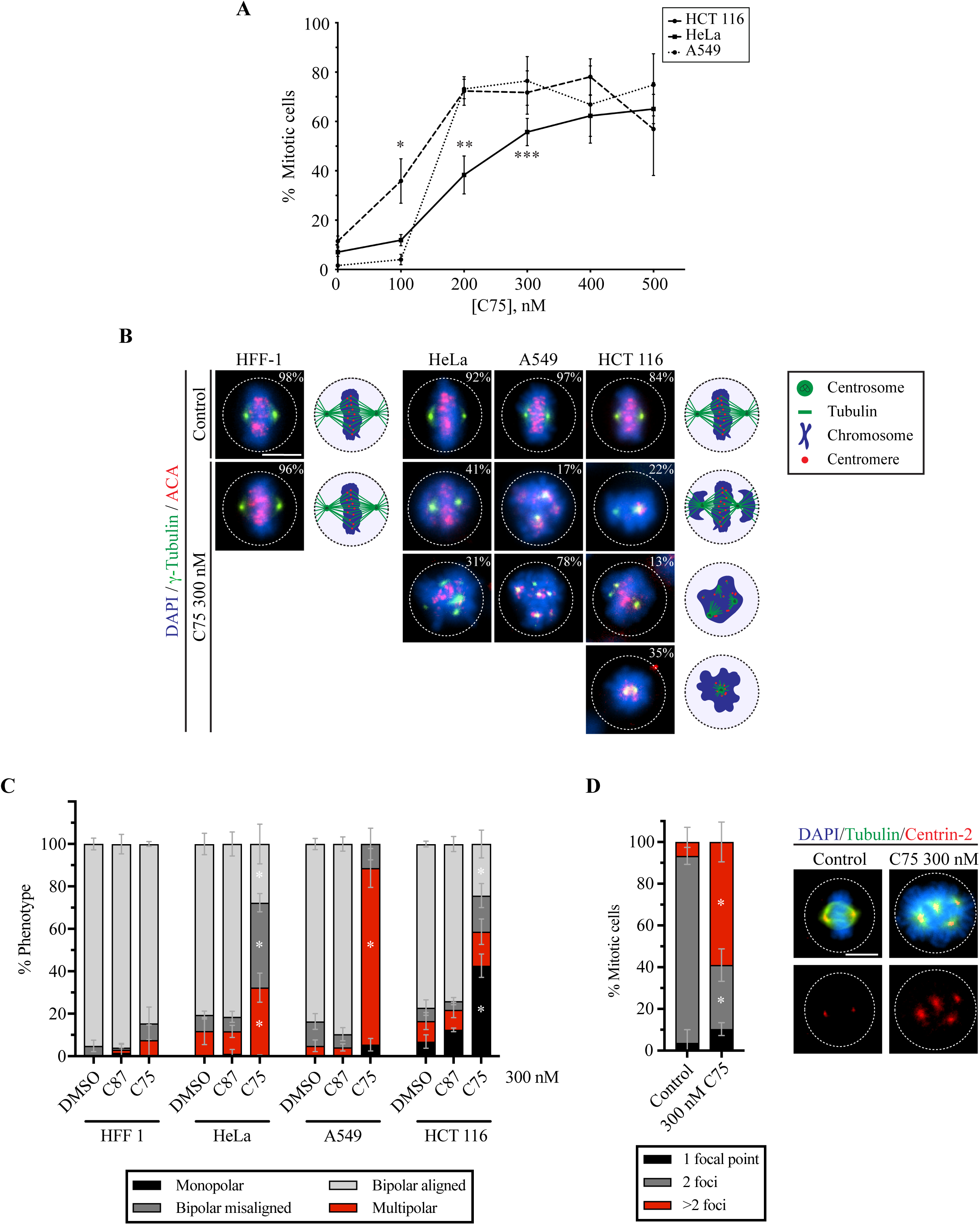
C75 causes multipolar spindles and mitotic arrest. **A)** A line graph shows changes in the proportion of mitotic HeLa, A549 and HCT 116 cells after treatment with increasing concentrations of C75 for 24 hours. Asterisks indicate statistical significance using two-way ANOVA test and post-hoc Tukey’s multiple comparison test with a 99% CI; a single asterisk means that HCT 116 is significantly different than HeLa and A549 (multiplicity adjusted *p* = 0.0007 and *p* < 0.0001, respectively), two indicates that HeLa is significantly different than A549 and HCT 116 (multiplicity adjusted *p* < 0.0001 for both), and three indicates that HeLa is significantly different than A549 (multiplicity adjusted *p* = 0.0035). **B)** Images show fixed HFF 1, HeLa, A549 and HCT116 cells stained for DNA (DAPI; blue), gamma-tubulin (green), and centromeres (ACA; red) after treatment with 300 nM of C75 for 4 hours. Cartoon schematics (green circles, centrosomes; blue, chromatin; green lines, microtubules; red, centromeres) show the different phenotypes observed, including bipolar spindles with aligned chromosomes (top), bipolar spindles with misaligned chromosomes (second from top), multipolar spindles (third from top), or monopolar spindles (bottom). The proportion of cells with each phenotype is shown in the top right corner of each image. The scale bar is 10 μm. **C)** A bar graph shows the proportion of each spindle phenotype (bipolar aligned in light grey; bipolar misaligned in dark grey; multipolar in red; monopolar in black) for HFF 1, HeLa, A549 and HCT 116 cells treated as in B. Bars show standard deviation and asterisks indicate two-way ANOVA test and a post-hoc Tukey’s multiple comparison test with a 99% CI, and multiplicity adjusted *p* =< 0.001 for each phenotype vs. DMSO and C87. **D)** A bar graph shows the proportion of HeLa cells with 1 (black), 2 (dark grey) or more (red) centrin-2 foci after treatment with DMSO or 300 nM C75. Cells were fixed and stained for DNA (DAPI; blue), tubulin (green), and centrin-2 (red). Asterisks indicate statistical significance of C75 with 2 foci and >2 foci compared to DMSO using Multiple t test with a 95% CI with a multiplicity adjusted *p* = 0.0009 and *p* = 0.0031, respectively. The scale bar is 10 μm.

**Figure 3.**
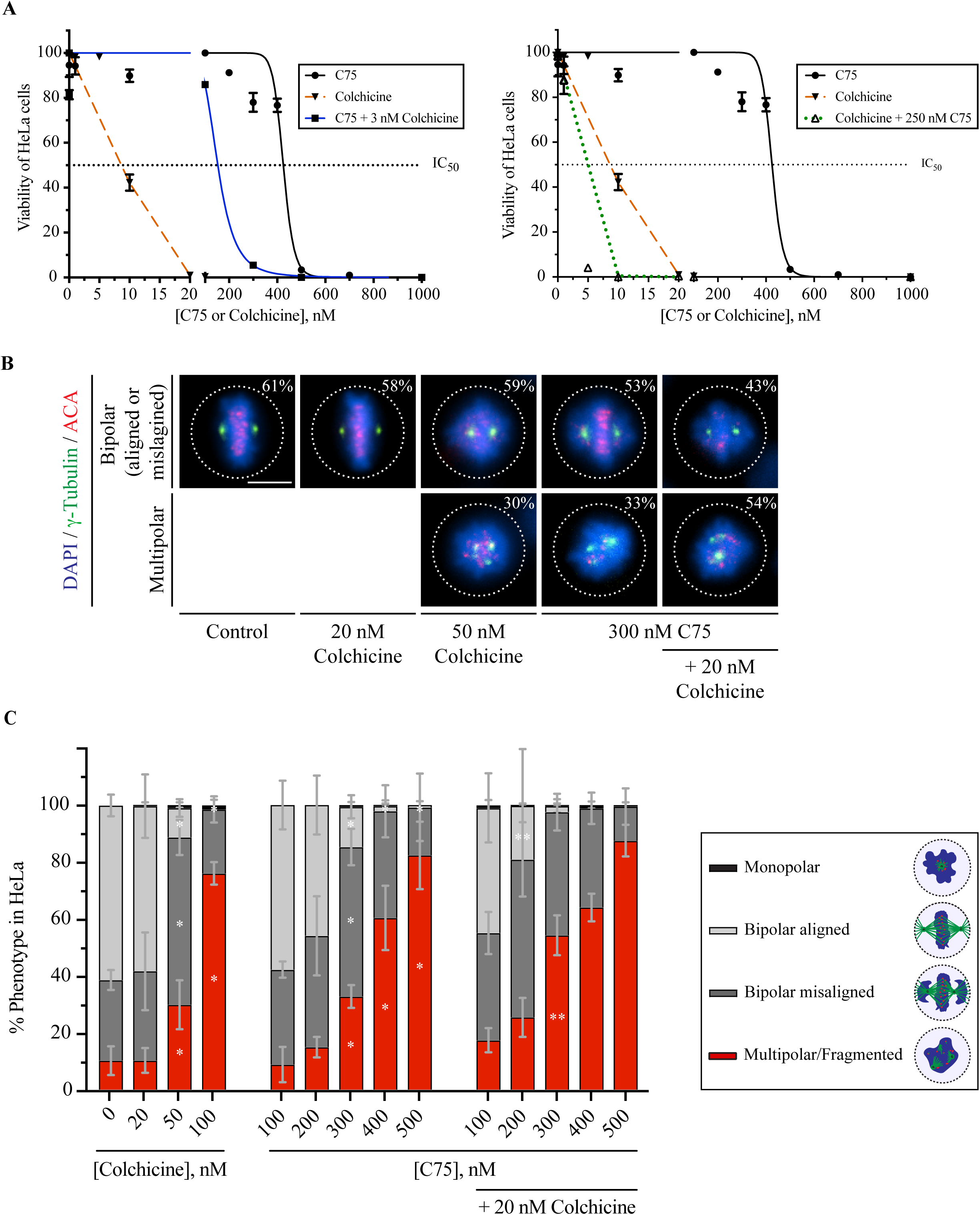
Combining C75 with Colchicine enhances lethality and spindle phenotypes. **A)** Line graphs show the IC_50_ for the viability (dotted black lines) of HeLa cells treated with varying concentrations of C75, Colchicine, or both after three population doubling times as indicated. The graph on the left shows changes in viability (Y-axis) with increasing C75 concentrations (X-axis; black line), Colchicine (dotted orange line), and C75 with 3 nM Colchicine (blue line). The graph on the right shows the changes in viability with Colchicine and 250 nM C75 (green dotted line). The bars indicate SEM, for N=3 experimental replicates. **B)** Images show fixed HeLa cells stained for DNA (DAPI; blue), gamma-tubulin (green) and ACA (centromeres; red) after treatment for 5 hours with Colchicine, C75 or both. On the images are the proportion of cells with bipolar spindles, and aligned chromosomes (control, DMSO or 20 nM Colchicine), or misaligned chromosomes and multipolar spindles (50 nM Colchicine, 300 nM C75 or both). The scale bar is 10 μm. **C)** Bar graphs show the proportion of HeLa cells from B) with bipolar aligned (light grey), bipolar misaligned (dark grey), multipolar (red) or monopolar (black) spindles. Bars show standard deviation. Statistical analysis was done using two-way ANOVA test and a post-hoc Tukey’s multiple comparison test with a 90% CI, and multiplicity adjusted *p* =< 0.0831; a single asterisk indicates significance to the Control (DMSO) and two indicates significance to C75.

**Figure 4.**
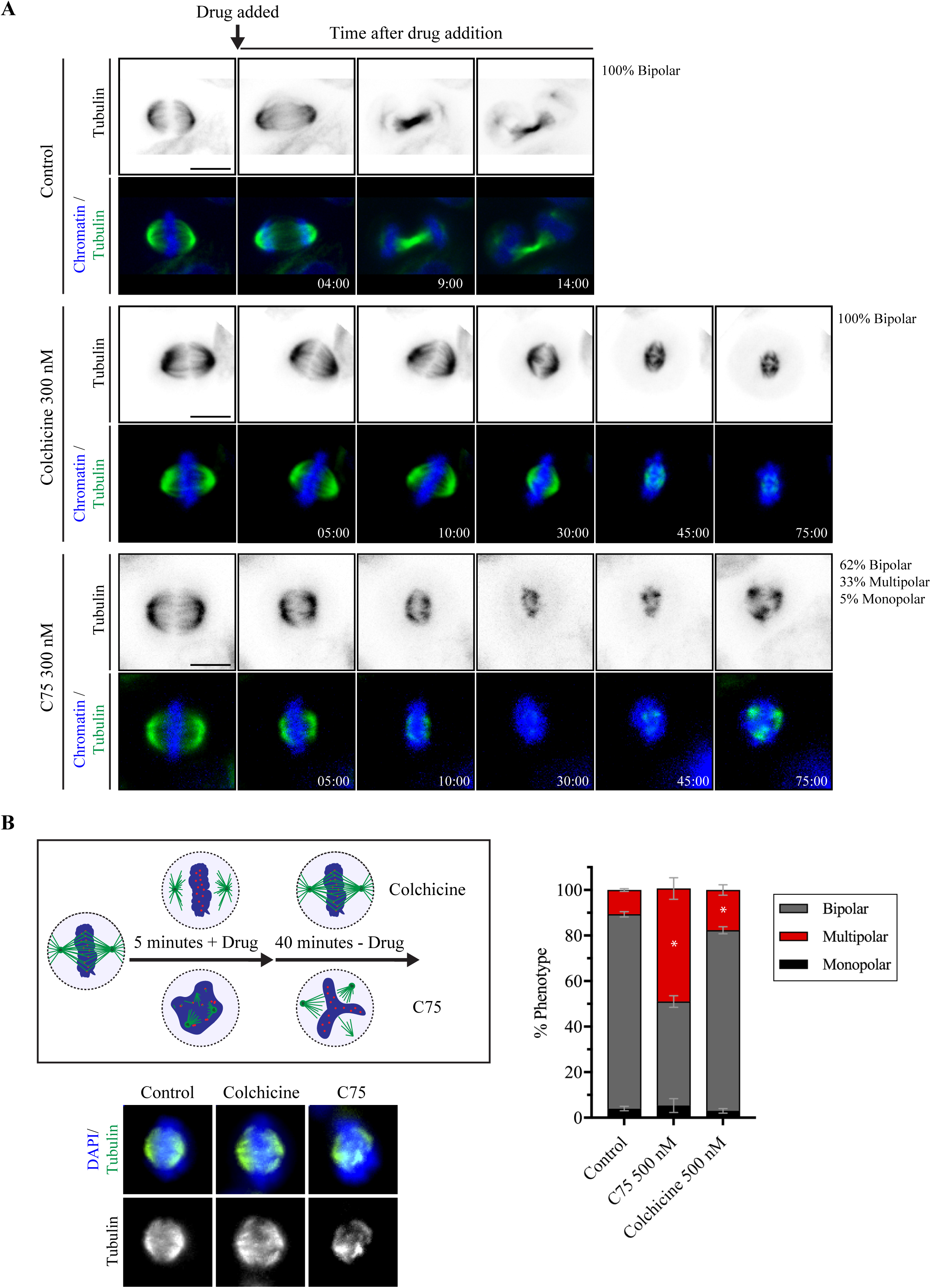
C75 and Colchicine cause different spindle phenotypes. **A)** Timelapse images show live HeLa cells co-stained for DNA (Hoechst 33342; blue) and microtubules (SiR-tubulin; green). Times are indicated in minutes. An arrow points to the time of addition of 300 nM of C75 (n=21), 300 nM of Colchicine (n=18) or DMSO (control; n=17). The proportion of cells with bipolar, multipolar or monopolar spindles are indicated. **B)** A cartoon schematic shows how HeLa cells were treated with drug (DMSO, 500 nM C75 or Colchicine) for 5 minutes, then fixed 40 minutes after the drugs were washed out. Underneath, images show fixed HeLa cells co-stained for DNA (DAPI; blue) and tubulin (green or white) after treatment with C75 or Colchicine. To the right, a bar graph shows the proportion of cells with monopolar (black), bipolar (dark grey) or multipolar (red) spindles. Bars show standard deviation and asterisks indicate *p*<0.05 by multiple t tests vs. control. The scale bar for all cells is 10 μm.

**Figure 5.**
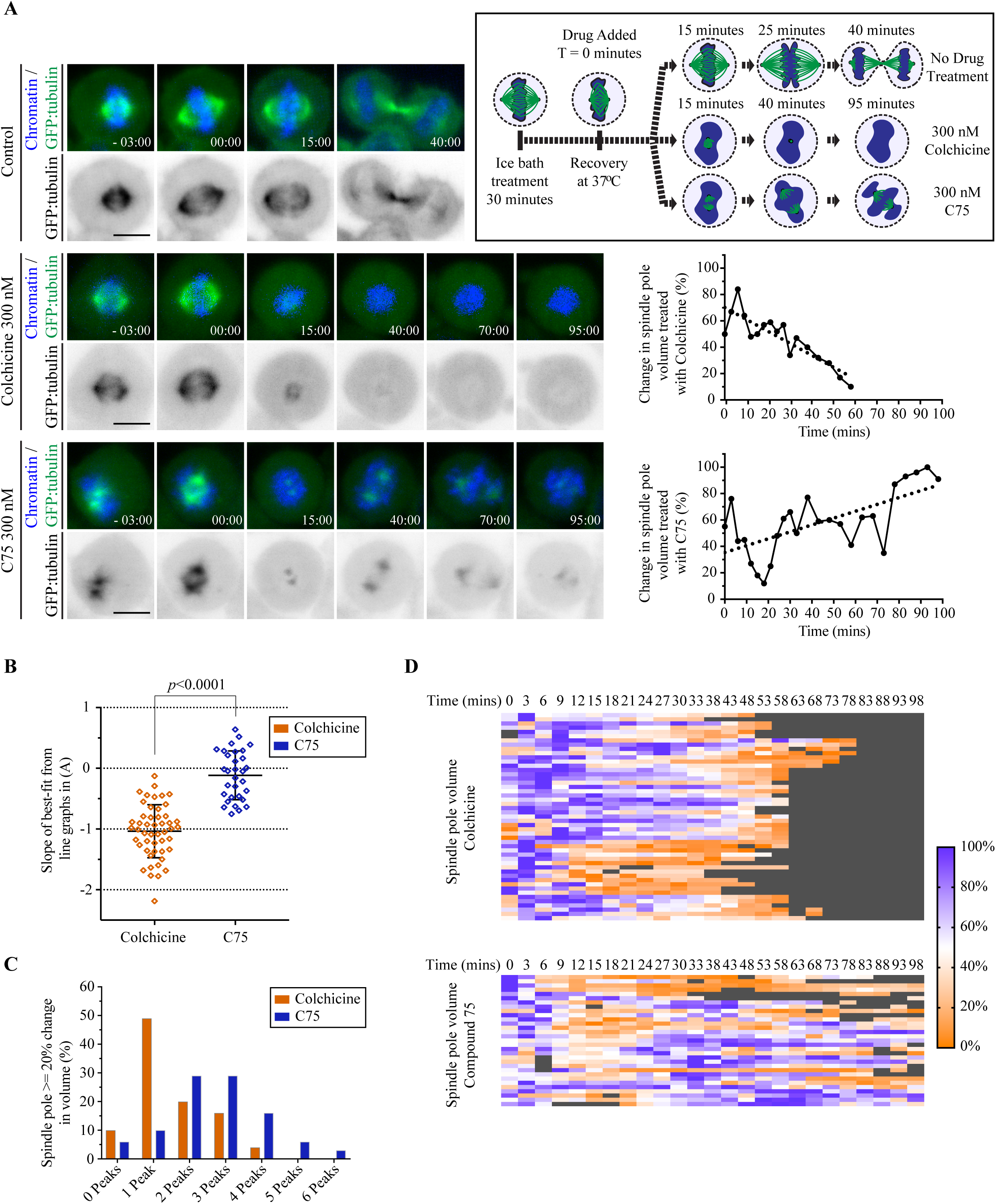
Microtubules and spindle poles recover in the presence of C75. **A)** A cartoon schematic (box at upper right) shows the experimental design. HeLa cells were cold-treated for 30 minutes in an ice bath to collapse microtubules, then upshifted to 37°C and imaged for microtubule regrowth in the presence or absence of 300 nM C75 or Colchicine. Timelapse images show live HeLa cells stably expressing GFP:tubulin (green) and co-stained for DNA by Hoechst (blue), with the times indicated in minutes. DMSO (control, n=6), Colchicine (n=24) or C75 (n=15) was added at 0 minutes. To the right, line graphs show changes in spindle pole volume (%; Y-axis) over time (X-axis) after addition of Colchicine or C75 (dotted line indicates best-fit). **B)** A scatter plot shows the distribution of slope values obtained from best-fit line graphs (Y-axis) for the changes in each spindle pole volume over time after treatment with C75 or Colchicine as in A). Significance was determined using the two-tailed Welch’s t test, *p***<**0.0001. **C)** A bar graph shows the number of peaks (X-axis) with an amplitude equal to or greater than 20% of the maximum volume of the spindle pole after treatment with C75 or Colchicine (Y-axis). **D)** Heat maps show the change in volume (%) of individual spindle poles over time (at the top in minutes) in cells treated with Colchicine or C75 as in A). Purple indicates larger volumes while orange indicates smaller volumes. Each line indicates a different spindle pole.

**Figure 6.**
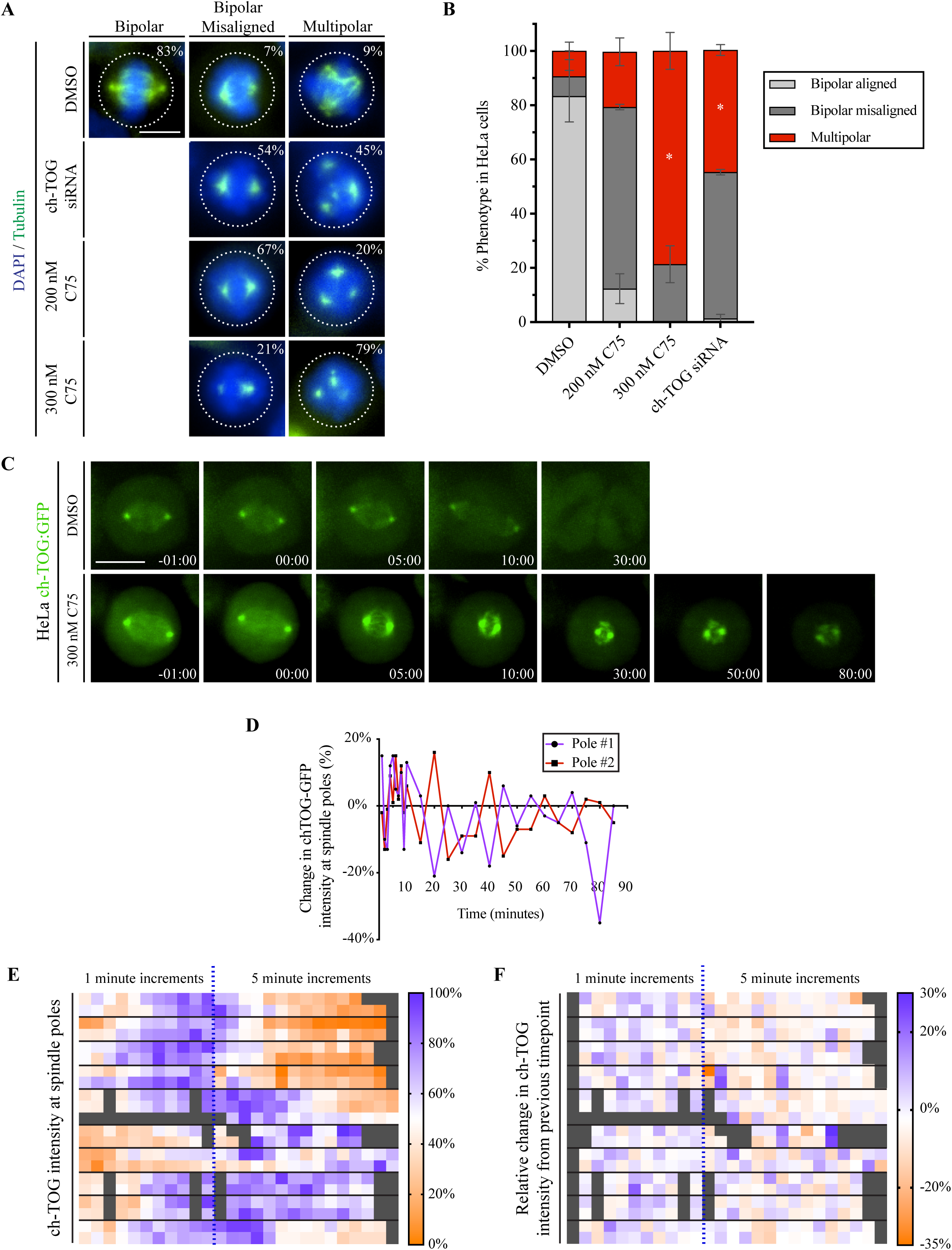
ch-TOG knockdown phenocopies C75, and ch-TOG oscillates in C75-treated cells. **A)** Images show fixed HeLa cells stained for DNA (DAPI, blue) and microtubules (tubulin, green) after treatment with ch-TOG siRNAs or 4 hours with C75 as indicated. The proportion of cells with bipolar spindles and aligned chromosomes, bipolar spindles with misaligned chromosomes, or multipolar spindles is indicated on the images. The scale bar is 10 μm. **B)** A bar graph shows the proportion of cells (%) with bipolar spindles (black), bipolar spindles with misaligned chromosomes (dark grey) or multipolar spindles (red) as in A. Bars show standard deviation. Asterisks show *p*<0.0001 with respect to Control (DMSO) using two-way ANOVA with a 99% confidence interval. **C)** Timelapse images show live HeLa cells expressing ch-TOG:GFP (green) in the presence (n=10) or absence of C75 (n=13). Times are indicated in minutes. The scale bar for all cells is 10 μm. **D)** A line graph shows the change in ch-TOG:GFP signal intensity (%, Y-axis) over time (minutes, X-axis) for each pole in an individual cell after C75 treatment as in C). **E)** A heat map show the change in intensity of ch-TOG:GFP (%) at the spindle poles. Times are indicated above, with timepoints from 1-11 in increments of 1 minute, and the remainder in increments of 5 minutes (separated by the blue dotted line). **F)** A heat map shows the relative change in intensity between two adjacent timepoints (%) after C75-treatment. Times are indicated above, with the same scale as in E). For both heat maps, purple indicates larger values, while orange indicates smaller values. The poles from the same cell are plotted together, and separated by cells via lines.

To determine synergy for the viability of cells treated with C75 and Colchicine in combination, CompuSyn software was used (Figure 3, Figure S3A). We used the non-constant ratio method of analysis since the concentration used for the combination studies was selected based on the highest concentration without lethality. The analysis indicated what combination of drug concentrations yielded an antagonistic, synergistic and additive effect based on the well-documented combination index (CI) described by the Chou-Talalay method (Chou 2010).

To determine the effect of C75 on spindle poles, HeLa cells stably expressing GFP:Tubulin and or ch-TOG:GFP; sh ch-TOG were used to measure changes in spindle volume and maximum intensity after upshift from cold treatment to recover depolymerized microtubules in the presence of C75 or Colchicine. To do this, we used the spot analyzer function in Imaris 9.5.1 (Bitplane), with parameters set to measure local contrast between the spindle pole and surrounding cytosol, which defined the pole boundaries (Supplemental Movies 1-3). Values were exported and organized in csv format using a macro in Python 3.0 to extract the volume and maximum intensity of each pole. The csv files were then imported into GraphPad Prism 7 to build graphical representations, including heat maps, bar graphs and distribution plots. Statistical analyses were also done in GraphPad Prism 7, to determine the significance between slope distributions using the Welch’s two-tailed t test with a 99% confidence interval (*p*<0.0001). Peaks corresponding to changes in spindle pole volume was determined for each pole using the built-in function “findpeaks” in MatLab R2017b. The parameters were set to identify any peaks that corresponded to a change in volume with an amplitude >= 20%. Identified peaks were then tabulated in Excel (Microsoft) and imported to GraphPad Prism 7 for graphical representation.

## Results

### Compound 75 causes cells to arrest in G2/M phase

We recently synthesized a new family of thienoisoquinoline compounds and identified several active derivatives that can bind to tubulin *in vitro* and cause reduced cell viability (*manuscript submitted*). The structure of Compound 75 (C75) is shown in Figure 1A, along with an inactive derivative C87. C75 prevents microtubule polymerization *in vitro* at concentrations equal to or above 250 nM (*e*.*g*. Fig. 1B) and disrupts previously assembled microtubule polymers (Fig. S1A). These characteristics of C75 are similar to Colchicine, which prevents polymerization *in vitro* at ranges of 1-5 μM, and depolymerizes microtubules (Fig. S1A; Fitzgerald 1976). C75 also binds to the Colchicine-pocket, as it can compete with Colchicine for binding to this site (Fig. S1B; Fitzgerald 1976; Hastie 1991). Together these data suggest that C75 binds to the Colchicine-binding site and prevents polymerization and destabilizes microtubules similar to Colchicine. However, C75 blocks polymerization at lower concentrations compared to Colchicine, suggesting that there are some differences in their accessibility and how they affect microtubules.

Next, we determined if C75 causes mitotic arrest similar to other tubulin-targeting compounds. While the inactive derivative, C87, had little effect on the viability of HFF-1 (human foreskin fibroblast), HeLa (cervical adenocarcinoma), A549 (lung carcinoma), or HCT 116 (colorectal carcinoma) cells, C75 caused reduced viability that varied depending on the cell line (Fig. 1C). While the IC_50_ for the viability of HeLa, A549 and HCT 116 cells treated with C75 was 427, 377 and 431 nM, respectively, it was 789 nM for HFF-1 cells (Fig. 1C). This suggests that cancer cells have ∼2-fold higher sensitivity to C75 compared to fibroblasts. To determine if the reduced viability is caused by cell cycle arrest, we performed flow cytometry on HeLa, A549 and HCT 116 cells after treatment with increasing concentrations of C75 for 8 hours. For all three lines, there was an increase in the proportion of cells in G2/M, and a decrease in G0/G1 (Figs 1D, S2). An ANOVA test followed by a post hoc Tukey’s multiple comparison test revealed that there was a significant change in the proportion of HeLa and A549 cells in G2/M and G0/G1 after treatment with 400 or 500 nM of C75 (Fig. 1D). HCT 116 cells responded at lower concentrations, with significant changes observed after treatment with 200 nM, 300, 400 and 500 nM of C75 (Fig. 1D). While there was no change in the proportion of HeLa or A549 cells in S phase, there was a decrease in HCT 116 cells in S phase after treatment with 500 nM of C75 (Fig. 1D). To further demonstrate that cells had mitotic phenotypes, cells were fixed and stained for DNA, microtubules and centromeres after treatment with 300 or 500 nM C75 and compared to control cells. Indeed, while metaphase HeLa, A549 and HCT 116 cells had bipolar spindles and aligned chromosomes, the spindles in treated cells had reduced microtubule intensity, misaligned chromosomes and fragmented spindle poles (Fig. 1E). Next, we counted the number of rounded, mitotic HeLa, A549 and HCT 116 cells after treatment with varying concentrations of C75 for 24 hours (Fig. 2A). For all cell lines, we saw a dramatic increase in the proportion of mitotic cells at 300 nM (55.7% HeLa, 76.4% A549 and 71.7% HCT 116, respectively). From these data we conclude that C75 is a tubulin-targeting compound that causes mitotic arrest due to aberrant mitotic spindles.

### C75 causes spindle pole phenotypes

As described above, C75 causes spindle phenotypes that increase in severity with concentration. These phenotypes range from reduced microtubules and misaligned chromosomes to fragmented spindle poles. To further quantify the spindle phenotypes caused by C75, HFF-1, HeLa, A549 and HCT 116 cells were treated with 300 nM C75 for 4 hours and stained for DNA, γ-tubulin and centromeres (Fig. 2B). Cells were treated at a lower vs. higher concentration of C75 and for a shorter period of time (4 hours vs. 8 or 24 hours) to capture a broader range of phenotypes. While HFF-1 cells had no obvious change in their spindle morphology, HeLa, A549 and HCT 116 cells showed spindle phenotypes that we classified as monopolar (centrosomes failed to separate), bipolar misaligned (misaligned sister chromatids) and multipolar spindles (fragmented or de-clustered centrosomes; Figs 2B, C). As determined by multiple t tests, there was a significant increase in HeLa cells with bipolar misaligned (41.1 vs. 7.1% control) and multipolar spindles (30.6 vs. 7.6% control), while there was a significant increase in A549 cells with monopolar (5 vs. 0% control) and multipolar spindles (77.6 vs. 3% control; Fig. 2C). There was a significant increase in HCT 116 cells with monopolar (34.5 vs. 9.5% control), bipolar misaligned (21.5 vs. 4.4% control) and multipolar spindles (12.5 vs. 6.9% control; Fig. 2C). To determine if multipolar spindles arose due to γ-tubulin displacement, or due to disruption of more core centrosome components, we repeated this experiment with HeLa cells and stained for DNA, tubulin and centrin-2 (Fig. 2D). Indeed, a significantly greater proportion of C75-treated cells had more than 1 or 2 centrin-2 foci compared to control cells (59 vs. 10%). Therefore, C75 is capable of displacing centrosome components. The different proportions of spindle phenotypes caused by C75 in the different cell lines likely reflects underlying differences in their genetic backgrounds and overall sensitivity to spindle disruption.

### Combining C75 and Colchicine enhance spindle phenotypes

Our data shows that C75 has characteristics similar to Colchicine and can bind to the Colchicine-pocket on tubulin. However, C75 is more effective at preventing polymerization *in vitro* and it could bind to alternate sites on tubulin. In addition, the two compounds could have different accessibility, or targets in cells. If C75 binds to a different site or target compared to Colchicine, they could have synergistic vs. additive effects when combined. We found that the IC_50_ for viability of C75 and Colchicine in HeLa cells was 425 nM and 9.5 nM, respectively (Fig. 3A). Adding 3 nM of Colchicine, a sub-optimal dose that caused little change in viability, to varying concentrations of C75 lowered the IC_50_ for C75 to 153 nM (Fig. 3A). Similarly, adding 250 nM of C75, which had a minimal impact on viability, to a range of Colchicine concentrations lowered the IC_50_ for Colchicine to 1.9 nM (Fig. 3A). To ascertain whether the combinatorial treatments were synergistic, additive or antagonistic, we analyzed the data using software for drug combination studies called CompuSyn (Chou 2010). Using the non-constant ratio method, we determined the Combination Index (CI) for each drug combination and found that C75 and Colchicine synergized at concentrations where they typically had low lethality (Fig. S3A). In particular, synergy was observed with 3 nM of colchicine and 300 nM of C75, and when 250 nM C75 was combined with 5 nM Colchicine, with CI values of 0.20 and 0.25, respectively (Fig. S3A).

Next, we determined the spindle phenotypes caused by combining the two compounds. HeLa cells were treated for 4 hours with 20 or 50 nM Colchicine, or 100, 200 or 300 nM of C75 with or without 20 nM Colchicine. Cells were fixed and stained for DNA, γ-tubulin and centromeres, and the proportion of cells with bipolar, bipolar misaligned or multipolar spindles was counted for each treatment (Fig. 3B, C). While control (DMSO) and cells treated with 20 nM Colchicine, or 100 nM or 200 nM C75 had similar proportions of bipolar misaligned or multipolar spindles, higher concentrations of Colchicine (50 nM) or C75 (300 nM) caused an increase in the proportion of cells with spindle phenotypes (Fig. 3B, C). When 20 nM of Colchicine was added to C75 (100, 200 and 300 nM) the proportion of cells with bipolar misaligned and/or multipolar spindles increased (Fig. 3B, C). In particular, combining 20 nM Colchicine with 300 nM C75 caused the multipolar phenotype to increase beyond the sum of each drug on their own at those concentrations. A similar enhancement of phenotypes was observed in HCT 116 cells, when 20 nM Colchicine was added with 300 or 400 nM of C75 (Fig. S3B). Therefore, C75 and Colchicine enhance each other’s effect in cells, suggesting that they have different binding sites for tubulin, or bind to different targets in cells.

### C75 and Colchicine have different effects on microtubule polymers in cells

Our data shows that C75 causes mitotic spindle phenotypes, which includes the formation of multipolar spindles. Although C75 acts similarly to Colchicine *in vitro*, their enhancement in cells suggests that there are differences in their mechanism of action. We characterized spindle phenotypes immediately after adding C75 and compared these to Colchicine. Live metaphase HeLa cells were treated with 300 nM C75 or Colchicine and imaged using live dyes (Hoechst 33342 to visualize chromatin, and SiR-tubulin to visualize tubulin) to monitor spindle phenotypes. The majority of control cells (DMSO-treated) exited mitosis, with bipolar spindles that matured to segregate sister chromatids and form a midbody (14/15), with the exception of one cell where a tripolar spindle formed into a pseudo-bipolar spindle and divided successfully (Fig. 4A). All cells treated with 300 nM Colchicine remained bipolar, and some cells arrested with collapsed spindles (5/20), while others exited mitosis similar to control (15/20; Fig. 4A). Cells treated with 300 nM C75 either arrested in mitosis with bipolar spindles (12/20), or became multipolar (7/20), and a monopolar spindle formed in one cell (1/20; Fig. 4A). In cells treated with either Colchicine or C75, the microtubules decreased or were lost altogether, but in C75-treated cells, microtubules re-polymerized with an uneven distribution between the two poles (Fig. 4A).

Next, we compared how cells recovered from short treatments of C75 or Colchicine. HeLa cells were treated with 500 nM of C75 or Colchicine for 5 minutes, then the drugs were washed out, and cells were left to recover in drug-free media for 40 minutes prior to fixing and staining them for DNA and tubulin (Fig. 4B). While the majority of cells recovered with bipolar spindles after Colchicine treatment (79.2% bipolar vs. 17.7% multipolar), a significantly larger proportion of cells treated with C75 had multipolar spindles determined by the multiple t test (45.7% bipolar vs. 49.7% multipolar; Fig. 4B). This data shows that C75 causes different spindle phenotypes compared to Colchicine in cells, although both compounds can prevent microtubule polymerization and cause depolymerization *in vitro*.

To further compare the effect of C75 and Colchicine on microtubules in cells, we induced microtubule depolymerization by cold treatment, then monitored their re-growth in the presence of the compounds. To do this we used HeLa cells stably expressing GFP-tagged tubulin, and cold-treated the cells for 30 minutes in an ice bath. The cells were then upshifted to 37°C at which time 300 nM C75 or Colchicine was added, then the cells were imaged during recovery as shown in Figure 5A. All of the control cells re-formed robust bipolar spindles and were in telophase ∼40 minutes after temperature upshift, indicating that the microtubules recovered and formed a functional mitotic spindle. In the presence of Colchicine, microtubules steadily decreased in intensity, with no detectable GFP signal by 98 minutes (24/24 cells; Fig. 5A). In C75-treated cells, microtubules similarly decreased, but after 20 minutes they polymerized and a GFP signal was detected in all cells (15/15 cells; Fig. 5A). The spindles were disorganized with misaligned sister chromatids, and cells failed to exit mitosis. To further characterize these phenotypes, we quantified changes in spindle pole volume and maximum intensity over time (Figs 5A-D, S4A, B). As expected, Colchicine-treatment caused a decrease in both volume and maximum intensity of the spindle poles over time (Figs 5A-D; S4A, B). Some fluctuation in volume was observed, however, there was a net linear decrease in volume and signal intensity over time as indicated by the best-line fits with negative slopes (Figs 5A, B; S4A, B). Heatmaps show the proportional changes in spindle volume or intensity for each spindle pole, and highlight how each pole decreased over time, albeit some sooner than others (Figs 5D, S4B, Supplemental Movie 1). Unlike Colchicine, C75-treatment caused an initial decrease in spindle pole volume and maximum intensity, followed by subsequent recovery and oscillation (Figs 5A, B; S4A, B). This is more evident in the heatmaps for each spindle pole, where signal intensity and pole volume continued to recover and fluctuate over time (Figs 5D, S4B, Supplemental Movie 2). To better understand this oscillatory pattern in spindle pole volume, we quantified the number of peaks where there was a >20% change in volume (Fig. 5C). While the majority of spindle poles in Colchicine-treated cells had only 1 or 2 peaks (69% with 1 or 2 vs. 31% with >2), the majority of poles in C75-treated cells had >2 peaks (46% with 1 or 2 vs. 54% with >2; Fig. 5C). Thus, C75 causes distinct phenotypes compared to Colchicine, where microtubule growth can recover in the presence of the compound, but this growth is not even between the spindle poles and leads to the formation of disorganized mitotic spindles.

Depleting ch-TOG, a microtubule polymerase, caused phenotypes similar to C75 (Fig. 6A; Barr and Bakal 2015; Gergely et al. 2003). HeLa cells treated with ch-TOG siRNAs had an increase in the proportion of bipolar spindles with misaligned chromosomes (51 vs. 17% for control) and multipolar or fragmented spindles (42 vs. 8% for control; Fig. 6A, B). As described earlier, we saw an increase in both phenotypes in cells treated with a mild increase in C75, from 200 to 300 nM C75 for 4 hours (64 and 22% bipolar misaligned; 24 and 76% multipolar, respectively; Fig. 6A, B). Since ch-TOG balances the activity of MCAK, a microtubule depolymerase to control microtubule length, we propose that C75 similarly causes excess microtubule depolymerization (Barr and Bakal 2015; Holmfeldt et al. 2004).

Next, we determined how C75 effects ch-TOG localization. Metaphase HeLa cells co-expressing sh ch-TOG for endogenous knockdown and sh-resistant ch-TOG:GFP were imaged immediately after treatment with 300 nM C75. In control cells, ch-TOG localized to the spindle poles and spindle microtubules, which decreased and was no longer detectable as cells progressed through mitotic exit (*e*.*g*. 30 minutes;13/13; Fig. 6C). In cells treated with 300 nM C75, ch-TOG increased in intensity at the spindle poles, which moved closer together within 5 minutes, and remained visible for more than 50 minutes (10/10; Fig. 6C). We also observed the accumulation of ch-TOG at a third pole in one of the cells, albeit with lower intensity compared to the other poles (*e*.*g*. Fig. 6E; Supplemental Movie 3). Measuring the change in maximum intensity of ch-TOG at spindle poles within an individual C75-treated cell revealed differences in the signal at one pole relative to the other, which oscillated over time (Fig. 6D). We then measured the intensity of ch-TOG in multiple C75-treated cells, and plotted these values as heatmaps of their maximum values (%), where purple shows values closest to the maximum, and orange shows the lowest (Fig. 6E). While there was a trend for ch-TOG levels to recover to maximum levels within the first 10 minutes of imaging, we observed reciprocal changes in intensity between the poles as in Figure 6D. To show this more clearly, we made a heat map of the relative changes in intensity from one timepoint to the next, where the colour reflects the increase (purple) or decrease (orange; Fig. 6E). Indeed, for most time points, there were inverse changes in the intensity of ch-TOG at the spindle poles within each cell, which started almost immediately after the addition of C75. This suggests that although ch-TOG accumulates at spindle poles, some pools of ch-TOG are displaced and oscillate between the poles, supporting their instability.

## Discussion

Here we describe the mechanism of action of C75, a new microtubule-targeting drug, in cultured human cells. C75 can compete with Colchicine for the Colchicine-binding pocket on tubulin and prevents microtubule polymerization *in vitro* (Figs 1, S1). C75 causes reduced viability and mitotic arrest due to the formation of aberrant spindles similar to other microtubule-targeting drugs including Colchicine (Figs 1, 2, S2). However, C75 has a different effect on microtubules in cells compared to Colchicine (Figs 4-6). For example, in HeLa cells, Colchicine causes a decrease in microtubules followed by gradual collapse of the spindle poles (Fig. 4), and fragmentation of the poles is only observed after long periods of time (*e*.*g*. hours vs. minutes; Figs 3, 6). However, C75 causes a decrease in microtubules and spindle pole fragmentation within minutes, followed by microtubule polymerization (Fig. 4). We also observed that tubulin or ch-TOG at the spindle poles oscillate during recovery, leading to the formation of disorganized spindles (Figs 5, 6, S4). In support of their different spindle phenotypes, combining C75 and Colchicine synergistically reduce cell viability and enhance spindle phenotypes when used at low concentrations (Figs 3, S3).

Our findings shed some light on the mechanism of action of C75 and how it affects microtubules in cells. The spindle phenotypes are similar to those caused by the depletion of ch-TOG, which is a microtubule polymerase (Fig. 6; Gergely et al. 2003; Brouhard et al. 2008; Barr and Bakal 2015). Many MAPs work together to form functional, bipolar spindles (Petry 2016). The loss of ch-TOG presumably causes an imbalance in the activities of other enzymes, including the microtubule depolymerase MCAK. Functions for MCAK include destabilizing microtubule ends to correct kinetochore attachments, which is controlled by Aurora B kinase phosphorylation, and focusing the asters into spindle poles via Aurora A kinase phosphorylation (Andrews et al. 2004; Ems-Mcclung et al. 2013; Walczak et al. 2013; Zhang et al. 2008). ch-TOG-depleted cells have phenotypes consistent with these roles including disorganized spindle poles and misaligned sister chromatids (Gergely et al. 2003). We similarly observed the formation of disorganized spindles and misaligned chromosomes in C75-treated cells. Based on this we speculate that: 1) C75 binds to tubulin dimers in a way that causes depolymerization of microtubules, but has a high off-rate and new polymers can form due to local increases in the critical concentration of free tubulin, and/or 2) C75 disrupts the microtubule or tubulin-binding site(s) of ch-TOG, mimicking ch-TOG loss-of-function. ch-TOG has multiple TOG domains that associate with tubulin dimers to mediate their assembly into polymers, as well as a basic region that can mediate interactions with the microtubule lattice (Widlund et al. 2011; Brouhard et al. 2008). In support of this we observed the accumulation and oscillation of ch-TOG between spindle poles in C75-treated cells (Fig. 6), which could reflect the displacement of ch-TOG from microtubules or tubulin. In addition, the spindle phenotypes caused by C75 are restricted to metaphase, which could reflect its inhibition of an enzyme like ch-TOG with a similar window of requirement.

Another microtubule-targeting compound, Vinblastine, similarly has complex effects on microtubules which could contribute to its success as an anti-cancer drug. *In vitro*, Vinblastine causes depolymerization at minus ends, and prevents growth at the plus ends (Panda et al., 1996). In cells, 2 or 10 nM of Vinblastine treatment causes disengagement of the mother and daughter centrioles and altered ultra-architecture of centrioles, or reduced microtubule dynamics and multipolar spindles, respectively (Jordan et al., 1992; Levrier et al. 2017; Wendell et al., 1993). We propose that C75 shares some similarities with Vinblastine with an accessibility and/or binding kinetics that favors microtubule depolymerization at the minus ends. But this accessibility appears to be restricted to cells in metaphase, as cells in other stages of the cell cycle do not appear to be affected at similar concentrations. Vinblastine is one of the few microtubule-destabilizing drugs that has been successful in the clinic as an anti-cancer therapy. There are many factors that impact how well a compound functions *in vivo*, and it is difficult to know what concentrations are reached at the target cells (Hughes et al. 2011). The ability of Vinblastine to cause spindle phenotypes at very low concentrations without disrupting the microtubule polymer mass could be ideal for minimizing side-effects. While C75 has a high IC_50_ value by comparison (in the 300-400 nM range for HeLa, HCT 116 and A549 cells), it’s unique effect on microtubules and high selectivity for metaphase cells could make it an interesting compound to consider for *in vivo* use. Importantly, the design of our compound is such that the scaffold is amenable to modifications at three different sites, and we are constantly making new derivatives with improved IC_50_ values and solubility that would replace C75 as our lead (*manuscript submitted*).

C75 causes spindle pole fragmentation prior to the oscillation of tubulin and ch-TOG at spindle poles. Centrosome integrity has been shown to rely on a TACC3-Clathrin-ch-TOG complex, and many microtubule-targeting drugs lead to the fragmentation of spindle poles, although this typically occurs after an extended period of time likely partly due to mitotic delay (*e*.*g*. hours; Foraker et al. 2012; Jordan et al. 1992; Karki and Shuster 2017). As mentioned above, low concentrations of Vinblastine cause disengagement of mother-daughter centrioles. In addition, cancer cells with high aneuploidy and amplified centrosomes typically rely on centrosome clustering mechanisms to form bipolar spindles (Leber et al. 2010; Ogden et al. 2012; Sabat-Pospiech et al. 2019). C75 may promote de-clustering and loss of centrosome integrity as visualized by gamma-tubulin and centrin-2 staining, making it an attractive compound to explore for the treatment of cancers with high centrosome numbers (Sabat-Pospiech et al. 2019). Indeed, different cancer cell lines have different proportions of cells with multipolar spindles after C75 treatment, presumably due to different genetic changes in their background including those that could impact centrosome integrity, clustering and/or microtubule dynamics (Fig. 2). We saw an increase in the proportion of A549 and HeLa cells with multipolar spindles, even at lower concentrations, suggesting that they are susceptible to de-clustering or fragmentation. Interestingly, we saw an increase in the proportion of HCT 116 cells with monopolar spindles at lower concentrations. With increased expression of Aurora A kinase and ch-TOG, these cells have increased assembly rates that could suppress C75 at lower concentrations (Ertych et al., 2014). The tendency of spindle poles to move closer together was observed in other cell types, but only after extensive microtubule depolymerization. Identifying specific genetic changes and correlating this with amplified centrosomes, or altered microtubule dynamics or regulation could predict how cells respond to C75-treatment.

## References

Akhmanova, Anna, and Michel O. Steinmetz. 2015. “Control of Microtubule Organization and Dynamics: Two Ends in the Limelight.” Nature Reviews Molecular Cell Biology 16 (12): 711–26. https://doi.org/10.1038/nrm4084.

Al-Bassam, Jawdat, and Fred Chang. 2011. “Regulation of Microtubule Dynamics by TOG-Domain Proteins XMAP215/Dis1 and CLASP.” Trends in Cell Biology 21 (10): 604–14. https://doi.org/10.1016/j.tcb.2011.06.007.

Andrews, Paul D., Yulia Ovechkina, Nick Morrice, Michael Wagenbach, Karen Duncan, Linda Wordeman, and Jason R. Swedlow. 2004. “Aurora B Regulates MCAK at the Mitotic Centromere.” Developmental Cell 6 (2): 253–68. https://doi.org/10.1016/S1534-5807(04)00025-5.

Barr, A. R., and F. Gergely. 2008. “MCAK-Independent Functions of Ch-Tog/XMAP215 in Microtubule Plus-End Dynamics.” Molecular and Cellular Biology 28 (23): 7199–7211. https://doi.org/10.1128/mcb.01040-08.

Barr, Alexis R, and Chris Bakal. 2015. “A Sensitised RNAi Screen Reveals a Ch-TOG Genetic Interaction Network Required for Spindle Assembly.” Scientific Reports 5 (April): 10564. https://doi.org/10.1038/srep10564.

Borisy, G. G., and E. W. Taylor. 1967. “The Mechanism of Action of Colchicine. Colchicine Binding to Sea Urchin Eggs and the Mitotic Apparatus.” The Journal of Cell Biology 34 (2): 535–48. https://doi.org/10.1083/jcb.34.2.535.

Brito, D, and CL Rieder. 2010. “The Ability to Survive Mitosis in the Presence of Microtubule Poisons” 66 (8): 437–47. https://doi.org/10.1002/cm.20316.The.

Brouhard, Gary J., Stear, Jeffrey H., Noetzel, Tim L. 2008. “XMAP215 Is a Processive Microtubule Polymerase.” Growth (Lakeland) 23 (1): 1–7. https://doi.org/10.1038/jid.2014.371.

Brouhard, Gary J., and Luke M. Rice. 2018. “Microtubule Dynamics: An Interplay of Biochemistry and Mechanics.” Nature Reviews Molecular Cell Biology 19 (7): 451–63. https://doi.org/10.1038/s41580-018-0009-y.

Byrnes, Amy E., and Kevin C. Slep. 2017. “TOG–Tubulin Binding Specificity Promotes Microtubule Dynamics and Mitotic Spindle Formation.” The Journal of Cell Biology. http://jcb.rupress.org/content/early/2017/05/15/jcb.201610090.

Chen, Fei, Nicholas W.Y. Wong, and Pat Forgione. 2014. “One-Pot Tandem Palladium-Catalyzed Decarboxylative Cross-Coupling and C-H Activation Route to Thienoisoquinolines.” Advanced Synthesis and Catalysis 356 (8): 1725–30. https://doi.org/10.1002/adsc.201300924.

Chou, Ting Chao. 2010. “Drug Combination Studies and Their Synergy Quantification Using the Chou-Talalay Method.” Cancer Research 70 (2): 440–46. https://doi.org/10.1158/0008-5472.CAN-09-1947.

Cirillo, Luca, Monica Gotta, and Patrick Meraldi. 2017. Cell Division Machinery and Disease. Vol. 1002. https://doi.org/10.1007/978-3-319-57127-0.

Dominguez-Brauer, Carmen, Kelsie L. Thu, Jacqueline M. Mason, Heiko Blaser, Mark R. Bray, and Tak W. Mak. 2015. “Targeting Mitosis in Cancer: Emerging Strategies.” Molecular Cell 60 (4): 524–36. https://doi.org/10.1016/j.molcel.2015.11.006.

Ems-Mcclung, Stephanie C., Sarah G. Hainline, Jenna Devare, Hailing Zong, Shang Cai, Stephanie K. Carnes, Sidney L. Shaw, and Claire E. Walczak. 2013. “Aurora B Inhibits MCAK Activity through a Phosphoconformational Switch That Reduces Microtubule Association.” Current Biology 23 (24): 2491–99. https://doi.org/10.1016/j.cub.2013.10.054.

Field, Jessica J., Arun Kanakkanthara, and John H. Miller. 2015. “Microtubule-Targeting Agents Are Clinically Successful Due to Both Mitotic and Interphase Impairment of Microtubule Function.” Bioorganic and Medicinal Chemistry 22 (18): 5050–59. https://doi.org/10.1016/j.bmc.2014.02.035.

Fielding, AB, S Lim, K Montgomery, I Dobreva, and S Dedhar. 2011. “A Critical Role of Integrin-Linked Kinase, Ch-TOG and TACC3 in Centrosome Clustering in Cancer Cells.” Oncogene 30 (5): 521–34. https://doi.org/10.1038/onc.2010.431.

Fitzgerald, Thomas J. 1976. “Molecular Features of Colchicine Associated with Antimitotic Activity and Inhibition of Tubulin Polymerization.” Biochemical Pharmacology 25: 1383–87.

Foraker, Amy B., Stéphane M. Camus, Timothy M. Evans, Sophia R. Majeed, Chih Ying Chen, Sabrina B. Taner, Ivan R. Corrêa, Stephen J. Doxsey, and Frances M. Brodsky. 2012. “Clathrin Promotes Centrosome Integrity in Early Mitosis through Stabilization of Centrosomal Ch-TOG.” Journal of Cell Biology 198 (4): 591–605. https://doi.org/10.1083/jcb.201205116.

Gergely, Fanni, Viji M. Draviam, and Jordan W. Raff. 2003. “The Ch-TOG/XMAP215 Protein Is Essential for Spindle Pole Organization in Human Somatic Cells.” Genes and Development 17 (3): 336–41. https://doi.org/10.1101/gad.245603.

Gigant, Benoît, Chunguang Wang, Raimond B.G. Ravelli, Fanny Roussi, Michel O. Steinmetz, Patrick A. Curmi, André Sobel, and Marcel Knossow. 2005. “Structural Basis for the Regulation of Tubulin by Vinblastine.” Nature 435 (7041): 519–22. https://doi.org/10.1038/nature03566.

Goodson, Holly V, and Erin M Jonasson. 2018. “Microtubules and Microtubule-Associated Proteins.”

Hanahan, Douglas, and Robert A. Weinberg. 2011. “Hallmarks of Cancer: The next Generation.” Cell 144 (5): 646–74. https://doi.org/10.1016/j.cell.2011.02.013.

Hastie, Susan Bane. 1991. “Interactions of Colchicine with Tubulin.” Pharmacology and Therapeutics 51 (3): 377–401. https://doi.org/10.1016/0163-7258(91)90067-V.

Holmfeldt, Per, Sonja Stenmark, and Martin Gullberg. 2004. “Differential Functional Interplay of TOGp/XMAP215 and the KinI Kinesin MCAK during Interphase and Mitosis.” EMBO Journal 23 (3): 627–37. https://doi.org/10.1038/sj.emboj.7600076.

Hood, Fiona E., Samantha J. Williams, Selena G. Burgess, Mark W. Richards, Daniel Roth, Anne Straube, Mark Pfuhl, Richard Bayliss, and Stephen J. Royle. 2013. “Coordination of Adjacent Domains Mediates TACC3-Ch-TOG-Clathrin Assembly and Mitotic Spindle Binding.” Journal of Cell Biology 202 (3): 463–78. https://doi.org/10.1083/jcb.201211127.

Hughes, J. P., S. S. Rees, S. B. Kalindjian, and K. L. Philpott. 2011. “Principles of Early Drug Discovery.” British Journal of Pharmacology 162 (6): 1239–49. https://doi.org/10.1111/j.1476-5381.2010.01127.x.

Jordan, M. A., D. Thrower, and L. Wilson. 1992. “Effects of Vinblastine, Podophyllotoxin and Nocodazole on Mitotic Spindles. Implications for the Role of Microtubule Dynamics in Mitosis.” Journal of Cell Science 102 (3): 401–16.

Jordan, Mary, and Kathy Kamath. 2007. “How Do Microtubule-Targeted Drugs Work? An Overview.” Current Cancer Drug Targets 7 (8): 730–42. https://doi.org/10.2174/156800907783220417.

Karki, Menuka, Neda Keyhaninejad, and Charles B. Shuster. 2017. “Precocious Centriole Disengagement and Centrosome Fragmentation Induced by Mitotic Delay.” Nature Communications 8 (May): 1–12. https://doi.org/10.1038/ncomms15803.

Kavallaris, Maria. 2010. “Microtubules and Resistance to Tubulin-Binding Agents.” Nature Reviews Cancer 10 (3): 194–204. https://doi.org/10.1038/nrc2803.

Kozielski, Frank. 2015. “Kinesins and Cancer.” Kinesins and Cancer 12 (August): 1–271. https://doi.org/10.1007/978-94-017-9732-0.

Kumar, Ashok, Parduman R. Sharma, and Dilip M. Mondhe. 2016. “Potential Anticancer Role of Colchicine-Based Derivatives: An Overview.” Anti-Cancer Drugs 28 (3): 250–62. https://doi.org/10.1097/CAD.0000000000000464.

Kumar, N. 1981. “Taxol-Induced Polymerization of Purified Tubulin. Mechanism of Action.” Journal of Biological Chemistry 256 (20): 10435–41.

Leber, Blanka, Bettina Maier, Florian Fuchs, Jing Chi, Phillip Riffel, Simon Anderhub, Ludmila Wagner, et al. 2010. “Proteins Required for Centrosome Clustering in Cancer Cells.” Science Translational Medicine 2 (33): 33ra38. https://doi.org/10.1126/scitranslmed.3000915.

Levrier, Claire, Martin C. Sadowski, Anja Rockstroh, Brian Gabrielli, Maria Kavallaris, Melanie Lehman, Rohan A. Davis, and Colleen C. Nelson. 2017. “6α-Acetoxyanopterine: A Novel Structure Class of Mitotic Inhibitor Disrupting Microtubule Dynamics in Prostate Cancer Cells.” Molecular Cancer Therapeutics 16 (1): 3–15. https://doi.org/10.1158/1535-7163.MCT-16-0325.

Martin, Maud, and Anna Akhmanova. 2018. “Coming into Focus: Mechanisms of Microtubule Minus-End Organization.” Trends in Cell Biology 28 (7): 574–88. https://doi.org/10.1016/j.tcb.2018.02.011.

Martino, Emanuela, Giuseppe Casamassima, Sonia Castiglione, Edoardo Cellupica, Serena Pantalone, Francesca Papagni, Marta Rui, Angela Marika Siciliano, and Simona Collina. 2018. “Vinca Alkaloids and Analogues as Anti-Cancer Agents: Looking Back, Peering Ahead.” Bioorganic and Medicinal Chemistry Letters 28 (17): 2816–26. https://doi.org/10.1016/j.bmcl.2018.06.044.

Massarotti, Alberto, Antonio Coluccia, Romano Silvestri, Giovanni Sorba, and Andrea Brancale. 2012. “The Tubulin Colchicine Domain: A Molecular Modeling Perspective.” ChemMedChem 7 (1): 33–42. https://doi.org/10.1002/cmdc.201100361.

Musacchio, Andrea. 2015. “The Molecular Biology of Spindle Assembly Checkpoint Signaling Dynamics.” Current Biology 25 (20): R1002–18. https://doi.org/10.1016/j.cub.2015.08.051.

Oakley, B. R., V. Paolillo, and Y. Zheng. 2015. “γ-Tubulin Complexes in Microtubule Nucleation and Beyond.” Molecular Biology of the Cell 26 (17): 2957–62. https://doi.org/10.1091/mbc.E14-11-1514.

Ogden, a, P C G Rida, and R Aneja. 2012. “Let’s Huddle to Prevent a Muddle: Centrosome Declustering as an Attractive Anticancer Strategy.” Cell Death and Differentiation 19 (8): 1255–67. https://doi.org/10.1038/cdd.2012.61.

Parness, J., and S. B. Horwitz. 1981. “Taxol Binds to Polymerized Tubulin in Vitro.” Journal of Cell Biology 91 (2 I): 479–87. https://doi.org/10.1083/jcb.91.2.479.

Petry, Sabine. 2016. “Mechanisms of Mitotic Spindle Assembly Sabine.” Annu Rev Biochem. 85 (1): 659–83. https://doi.org/10.1146/annurev-biochem-060815-014528.

Sabat-Pospiech, Dorota, Kim Fabian-Kolpanowicz, Ian A. Prior, Judy M. Coulson, and Andrew B. Fielding. 2019. “Targeting Centrosome Amplification, an Achilles’ Heel of Cancer.” Biochemical Society Transactions 47 (5): 1209–22. https://doi.org/10.1042/BST20190034.

Walczak, Claire E., Sophia Gayek, and Ryoma Ohi. 2013. “Microtubule-Depolymerizing Kinesins.” Annual Review of Cell and Developmental Biology 29: 417–41. https://doi.org/10.1146/annurev-cellbio-101512-122345.

Wang, Yuxi, Hang Zhang, Benoît Gigant, Yamei Yu, Yangping Wu, Xiangzheng Chen, Qinhuai Lai, Zhaoya Yang, Qiang Chen, and Jinliang Yang. 2016. “Structures of a Diverse Set of Colchicine Binding Site Inhibitors in Complex with Tubulin Provide a Rationale for Drug Discovery.” FEBS Journal 283 (1): 102–11. https://doi.org/10.1111/febs.13555.

Weaver, Beth A. 2014. “How Taxol/Paclitaxel Kills Cancer Cells.” Molecular Biology of the Cell 25 (18): 2677–81. https://doi.org/10.1091/mbc.E14-04-0916.

Widlund, Per O., Jeffrey H. Stear, Andrei Pozniakovsky, Marija Zanic, Simone Reber, Gary J. Brouhard, Anthony A. Hyman, and Jonathon Howard. 2011. “XMAP215 Polymerase Activity Is Built by Combining Multiple Tubulin-Binding TOG Domains and a Basic Lattice-Binding Region.” Proceedings of the National Academy of Sciences of the United States of America 108 (7): 2741–46. https://doi.org/10.1073/pnas.1016498108.

Wordeman, Linda. 2019. “GTP-Tubulin Loves Microtubule plus Ends but Marries the Minus Ends.” Journal of Cell Biology 218 (9): 2822–23. https://doi.org/10.1083/jcb.201908039.

Yu, Jun Xiong, Qian Chen, Ya Qun Yu, Shu Qun Li, and Jian Fei Song. 2016. “Upregulation of Colonic and Hepatic Tumor Overexpressed Gene Is Significantly Associated with the Unfavorable Prognosis Marker of Human Hepatocellular Carcinoma.” American Journal of Cancer Research 6 (3): 690–700.

Zasadil, Lauren M, Kristen A Andersen, Dabin Yeum, Gabrielle B Rocque, Lee G Wilke, Amye J Tevaarwerk, Ronald T Raines, Mark E Burkard, and Beth A Weaver. 2014. “Cytotoxicity of Paclitaxel in Breast Cancer Is Due to Chromosome Missegregation on Multipolar Spindles.” Science Translational Medicine 6 (229): 229ra43. https://doi.org/10.1126/scitranslmed.3007965.

Zhang, Xin, Stephanie C. Ems-McClung, and Claire E. Walczak. 2008. “Aurora A Phosphorylates MCAK to Control Ran-Dependent Spindle Bipolarity.” Molecular Biology of the Cell 19 (July): 2752–65. https://doi.org/10.1091/mbc.e08-02-0198.

